# Self-supervised deep learning for pan-cancer mutation prediction from histopathology

**DOI:** 10.1101/2022.09.15.507455

**Authors:** Oliver Lester Saldanha, Chiara M. L. Loeffler, Jan Moritz Niehues, Marko van Treeck, Tobias P. Seraphin, Katherine Jane Hewitt, Didem Cifci, Gregory Patrick Veldhuizen, Siddhi Ramesh, Alexander T. Pearson, Jakob Nikolas Kather

## Abstract

The histopathological phenotype of tumors reflects the underlying genetic makeup. Deep learning can predict genetic alterations from tissue morphology, but it is unclear how well these predictions generalize to external datasets. Here, we present a deep learning pipeline based on self-supervised feature extraction which achieves a robust predictability of genetic alterations in two large multicentric datasets of seven tumor types.

## Main Text

The genotype of any solid tumor determines its phenotype, giving rise to a large variety of patterns in cancer histopathology. Deep learning (DL), a tool from the realm of artificial intelligence, can infer genetic alterations directly from routine histopathology slides stained with hematoxylin and eosin (H&E). Initial studies demonstrated this predictability in lung cancer^1^, breast cancer^2^ and colorectal cancer.^3^ Since then, several “pan-cancer” studies have shown that DL-based prediction of biomarkers is feasible across the whole spectrum of human cancer.^4–7^. However, these studies are overwhelmingly performed in a single large cohort without externally validating the results on a large scale. This raises a number of potential concerns, as prediction performance can be heavily biased by batch effects occurring in such single multicentric datasets.^8^ To move closer to clinical applicability, external validation of any DL system is paramount.^9^ Finally, more recent technical benchmark studies have demonstrated that attention-based multiple instance learning (MIL)^10^ and self-supervised pre-training of feature extractors^11,12^ improves prediction performance for biomarkers in computational pathology, but these technical advances have not yet been systematically applied to multi-cohort mutation prediction.

We acquired two large, multi-centric datasets of cancer histopathology images with matched genetic profiling: the Cancer Genome Atlas (TCGA) and the Clinical Proteomic Tumor Analysis Consortium (CPTAC) dataset. We used all tumor types which were present in both datasets, namely: breast (BRCA; TCGA N=1062, CPTAC N=198 patients), colorectal (CRC; TCGA N=615, CPTAC N=178), glioblastoma (GBM; TCGA N=388, CPTAC N=189), lung adeno (LUAD; TCGA N=478, CPTAC N=244), lung squamous (LUSC; TCGA N=478, CPTAC N=212), pancreatic (PAAD; TCGA N=183, CPTAC N=168 patients) and (uterine) endometrial cancer (UCEC; TCGA N=505, CPTAC N=250; **Figure 1A-B**; **Suppl. Figure 1A-B**). We compiled a list of clinically relevant and targetable mutations of N=1066 genes from OnkoKb^13^, as well as mutation data from https://www.cbioportal.org/^14^, as shown in https://github.com/KatherLab/cancer-metadata. We used genes with a minimum of N=25 mutants in TCGA, resulting in 321 analyzable genes in endometrial cancer (UCEC), down to 4 analyzable genes in pancreatic cancer (PAAD, **Figure 1C, Figure 2A**). We developed a fully automatic DL pipeline based on a self-supervised feature extractor (https://github.com/Xiyue-Wang/RetCCL) and attention-based multiple instance learning (attMIL)^15^. We trained models to predict mutations in TCGA (**Figure 1A**) and evaluated the performance on CPTAC (**Figure 1B**). We report the mean (±standard deviation) area under the receiver operating characteristic curve (AUROC) of five replicate experiments.

**Figure 1:**
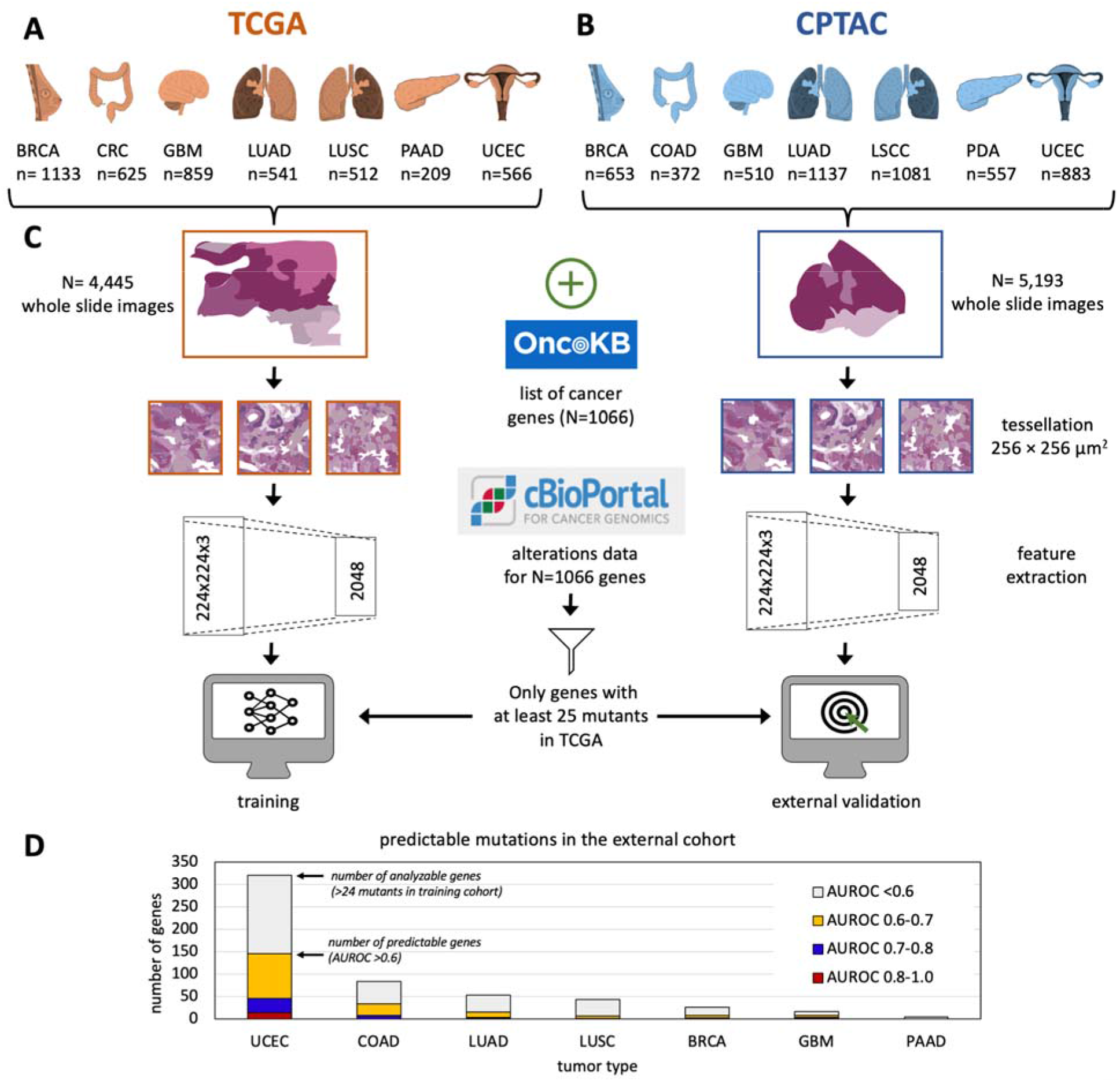
Prediction of mutations in TCGA and validation in CPTAC. **(A)** We acquired pathology images from The Cancer Genome Atlas (TCGA) **(B)** and the Clinical Proteomic Tumor Analysis Consortium (CPTAC). **(C)** We collated genetic mutations and trained Deep Learning classifiers to predict them from pathology slides in TCGA and externally validated them on CPTAC. (Icon source https://smart.servier.com/ www.flaticon.com, CC-BY). **(D)** Analyzable and predictable mutations in all cohorts, AUROC = area under the receiver operating characteristic curve.

**Figure 2:**
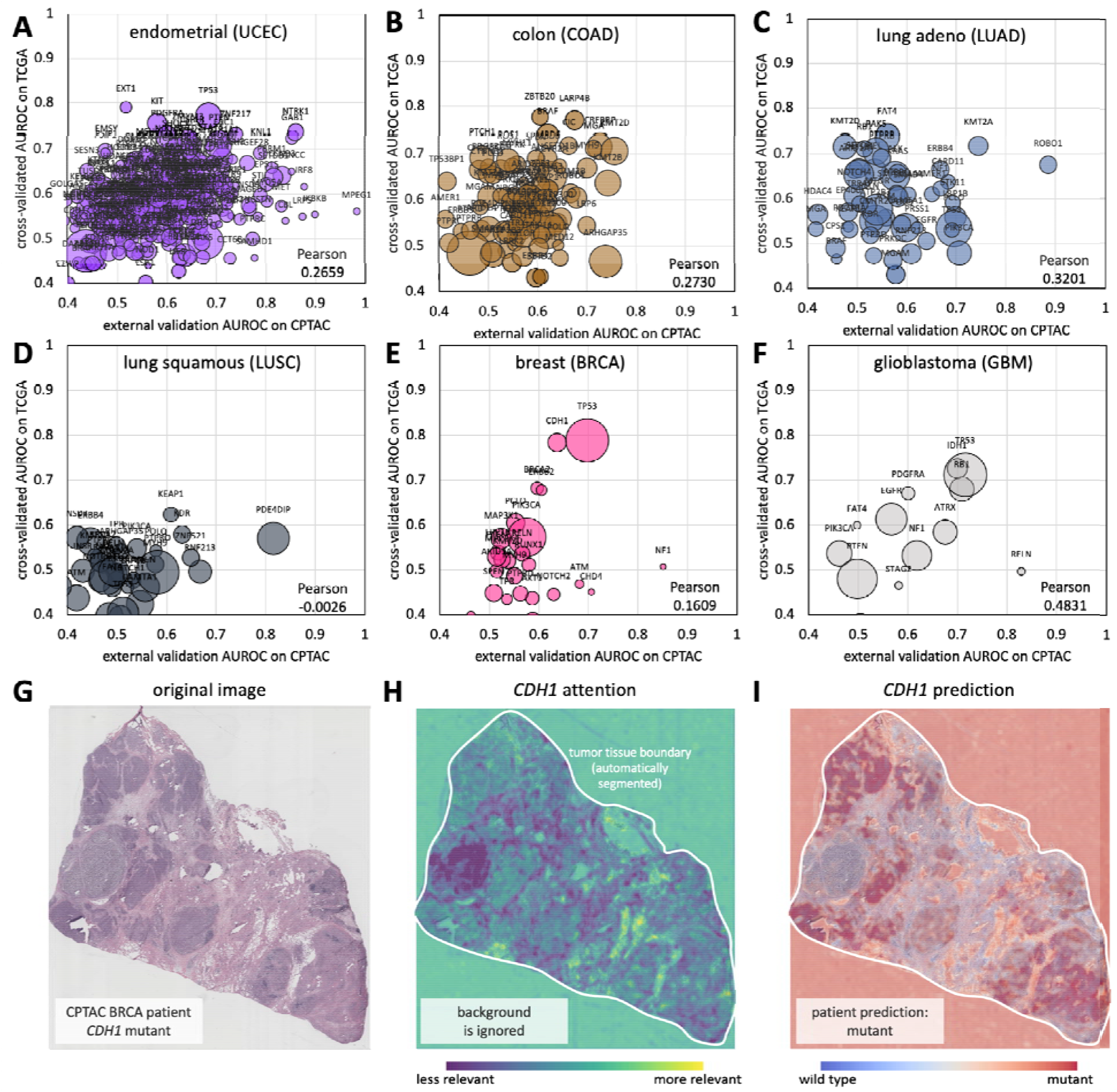
Classification performance for all genes in internal and external validation. **(A-F)** Internal cross-validated and external validation AUROC for six tumor types (PAAD in **Suppl. Figure 2**). The bubble size scales with the number of mutant patients in the external validation cohort. All raw data are in Suppl. Table 1. **(G-I)** A representative patient from the CPTAC-BRCA cohort, with attention map and prediction maps for *CDH1* mutational status.

In most tumor types, several genes were predictable from histology (**Figure 1D, Suppl. Table 1**). In accordance with previous studies^16^, endometrial cancer (UCEC) had the highest number of detectable mutations. N=145 out of n=321 analyzable genes had an with AUROC of over 0.60, of whom 31 had an AUROC between 0.70 and 0.80 in the external validation cohort, and an additional 14 genes had an AUROC over 0.80 (**Figure 2A**). Among these were *NTRK1* mutations (AUROC 0.86±0.07), which are potentially clinically relevant^17^; *MSH2* mutations (AUROC 0.73±0.26), which can cause microsatellite instability and are druggable with immune checkpoint inhibitors, and PTEN, which reached an AUROC of 0.71 (±0.05) and is involved in hereditary cancer.^18^ We identified 51 predictable genes (out of 84 analyzable genes) for colorectal cancer (CRC) with AUROCs of over 0.6, including 7 predictable genes with AUROCs over 0.7 in the external validation cohort. (**Figure 2B**). This included prognostic alterations, such as *CREBBP^19^* mutations, which reached an AUROC of 0.73±0.04 (**Suppl. Table 1**). Compared to the other tumor types in our study, the tumors with the most predictable genes (UCEC, COAD and LUAD) have a higher average tumor mutational burden.^20^ We hypothesize that many morphological alterations are related to immune-mediated changes in the tumor microenvironment. Our method yielded interpretable spatial predictions (**Figure 2G**), and unlike previous studies^21^ provided separate heatmaps for attention (**Figure 2H**) and classification (**Figure 2I, Suppl. Figure 3A-I**). A key limitation is that many clinically relevant genes were not analyzable due to having fewer than 25 mutants in TCGA. Large-scale efforts are needed to create datasets with a sufficient size, which could be facilitated by federated^22^ or swarm^23^ learning.

Since the early 2000s, studies have shown a link between genetic alterations and histological phenotypes^24^, which DL can exploit.^4,5^ While there is no biological reason why every frequent genetic alteration is actually manifest in histology, our results add to the growing amount of evidence which shows that many of these alterations are indeed determinable from H&E. Crucially, in contrast to previous studies, all our models have been externally validated, thereby minimizing the risk of overfitting.^8^ Our analysis shows that in tumor types with numerous predictable mutations, crossvalidated performance of attMIL is correlated to external validation performance (**Figure 2A-F, Supp. Figure 2**). Our study identifies several clinically relevant candidate genes amenable to DL-based prescreening as part of clinical routine practice, with the aim of identifying patients who are good candidates for confirmatory genetic testing.

## Additional Information

### Ethics statement

All experiments were conducted in accordance with the Declaration of Helsinki. For this study, we used anonymized H&E-stained slides from public repositories.

### Data availability

TCGA images are from https://portal.gdc.cancer.gov/, CPTAC images are from https://wiki.cancerimagingarchive.net/display/Public/CPTAC+Pathology+Slide+Downloads. Genetic data are available at https://www.cbioportal.org/.

### Code availability

Our pipeline is available under an open-source license (https://github.com/KatherLab/marugoto).

### Author contributions

OLS, CMML, SR, ATP and JNK conceived the experiments. MVT developed the codes for analysis. JMN tested and corrected the codes. CMMS and TPS curated the source data. KJH quality-checked the source data. DC and GPV quality-checked the genetic data. OLS performed the experiments. All authors interpreted the data and wrote the paper. All authors agreed to the submission of this paper.

## References

1. Coudray, N. et al. Classification and mutation prediction from non–small cell lung cancer histopathology images using deep learning. Nat. Med. 24, 1559–1567 (2018).

2. Couture, H. D. et al. Image analysis with deep learning to predict breast cancer grade, ER status, histologic subtype, and intrinsic subtype. NPJ Breast Cancer 4, 30 (2018).

3. Kather, J. N. et al. Deep learning can predict microsatellite instability directly from histology in gastrointestinal cancer. Nat. Med. 25, 1054–1056 (2019).

4. Fu, Y. et al. Pan-cancer computational histopathology reveals mutations, tumor composition and prognosis. Nature Cancer 1–11 (2020).

5. Kather, J. N. et al. Pan-cancer image-based detection of clinically actionable genetic alterations. Nature Cancer 1, 789–799 (2020).

6. Arslan, S. et al. Deep learning can predict multi-omic biomarkers from routine pathology images: A systematic large-scale study. bioRxiv 2022.01.21.477189 (2022) doi:10.1101/2022.01.21.477189.

7. Chen, R. J. et al. Pan-cancer integrative histology-genomic analysis via multimodal deep learning. Cancer Cell 40, 865–878.e6 (2022).

8. Howard, F. M. et al. The impact of site-specific digital histology signatures on deep learning model accuracy and bias. Nat. Commun. 12, 4423 (2021).

9. Kleppe, A. et al. Designing deep learning studies in cancer diagnostics. Nat. Rev. Cancer 21, 199–211 (2021).

10. Laleh, N. G. et al. Benchmarking weakly-supervised deep learning pipelines for whole slide classification in computational pathology. Med. Image Anal. 102474 (2022).

11. Schirris, Y., Gavves, E., Nederlof, I., Horlings, H. M. & Teuwen, J. DeepSMILE: Contrastive self-supervised pre-training benefits MSI and HRD classification directly from H&E whole-slide images in colorectal and breast cancer. Med. Image Anal. 102464 (2022).

12. Ciga, O., Xu, T. & Martel, A. L. Self supervised contrastive learning for digital histopathology. arXiv [eess.IV] (2020).

13. Chakravarty, D. et al. OncoKB: A Precision Oncology Knowledge Base. JCO Precis Oncol 2017, (2017).

14. Gao, J. et al. Integrative analysis of complex cancer genomics and clinical profiles using the cBioPortal. Sci. Signal. 6, l1 (2013).

15. Ilse, M., Tomczak, J. M. & Welling, M. Attention-based Deep Multiple Instance Learning. arXiv [cs.LG] (2018).

16. Loeffler, C. M. L. et al. Predicting Mutational Status of Driver and Suppressor Genes Directly from Histopathology With Deep Learning: A Systematic Study Across 23 Solid Tumor Types. Front. Genet. 12, (2022).

17. Hechtman, J. F. NTRK insights: best practices for pathologists. Mod. Pathol. 35, 298–305 (2022).

18. Spurdle, A. B., Bowman, M. A., Shamsani, J. & Kirk, J. Endometrial cancer gene panels: clinical diagnostic vs research germline DNA testing. Mod. Pathol. 30, 1048–1068 (2017).

19. Liu, K., Wang, J.-F., Zhan, Y., Kong, D.-L. & Wang, C. Prognosis model of colorectal cancer patients based on NOTCH3, KMT2C, and CREBBP mutations. J. Gastrointest. Oncol. 12, 79–88 (2021).

20. Chalmers, Z. R. et al. Analysis of 100,000 human cancer genomes reveals the landscape of tumor mutational burden. Genome Med. 9, 34 (2017).

21. Lu, M. Y. et al. Data-efficient and weakly supervised computational pathology on whole-slide images. Nature Biomedical Engineering 1–16 (2021).

22. Lu, M. Y. et al. Federated learning for computational pathology on gigapixel whole slide images. Med. Image Anal. 76, 102298 (2022).

23. Saldanha, O. L. et al. Swarm learning for decentralized artificial intelligence in cancer histopathology. Nat. Med. (2022) doi:10.1038/s41591-022-01768-5.

24. Rosner, A. et al. Pathway pathology: histological differences between ErbB/Ras and Wnt pathway transgenic mammary tumors. Am. J. Pathol. 161, 1087–1097 (2002).

